# A cytological framework of female meiosis in *Arabidopsis*

**DOI:** 10.1101/2025.08.22.671760

**Authors:** Bingyan Hu, Maria Ada Prusicki, Katharina Stahlmann, Yingqi Wang, Arp Schnittger

## Abstract

Female and male meiosis often differ in many aspects, such as their duration and the frequency as well as the positioning of crossovers. However, studying female meiosis is often very challenging and thus, much less is known about female versus male meiosis in many species including plants. To approach this gap, we have developed a live-cell imaging system for female meiocytes in Arabidopsis. This allowed us to obtain a temporally resolved cytological framework of female meiosis in the wildtype that serves as a guiding system for future studies. Here, we have applied this imaging system to study mutants in cyclin-dependent kinase inhibitors, in which a designated female meiocyte undergoes several mitotic divisions before entering meiosis. This enabled us to address when a meiocyte is committed to meiosis, a key question during reproductive development and in particular for the analysis of apomictic species in which meiosis is skipped.

**Highlights:** Establishment of a live-cell imaging system captures dynamic features of female meiosis. Identification of cytological landmarks ensures robust assignment of meiotic stages.

Time-lapse imaging enables quantitative dissection of meiotic phases. Application of the framework reveals great plasticity in the commitment to meiosis.

## Introduction

Meiosis is key for sexual reproduction and a major driving force for evolution. It ensures genetic stability across generations by reducing the genomic content by half so that the full genomic complement is restored after fertilization. In addition, meiosis promotes genetic diversity through reciprocal exchanges of genetic material between homologous chromosomes. The molecular machinery of meiotic recombination and chromosome segregation is highly conserved among such diverse eukaryotes as mammals and flowering plants, suggesting that meiosis was probably invented over one billion years ago in the last common ancestor. Given its fundamental characteristics and high degree of conservation, it is notable that there is a substantial level of dimorphism between the sexes in one species. One of the most prominent examples of the specifics of female meiosis is seen in mammals, where oocytes start meiosis synchronously together but get arrested twice during meiosis, the prophase I arrest lasting for years and even decades. In contrast, spermatocytes in males continuously undergo meiosis without interruption.^1,2^ Female meiosis yields only one viable gamete compared to four in males, and human oocytes show higher crossover (CO) frequencies than spermatocytes.^3–5^ Similarly, flowering plants show differences between female and male meiosis. For instance, the CO frequency is lower in female meiocytes (megaspore mother cells, MMCs) than in male meiocytes (pollen mother cells, PMCs) in *Arabidopsis thaliana* (Arabidopsis).^6,7^ Mutants in meiotic regulators such as the A-type cyclin TARDY ASYNCHRONOUS MEIOSIS or the anaphase-promoting complex inhibitor OMISSION OF SECOND MEIOTIC DIVISION/GIGAS CELLS, affect female versus male meiosis to a different level, indicating that the full extent and the molecular basis of these differences are far from being understood.^8,9^

Over the past decades, more than 100 plant genes have been functionally studied in the regulation of meiosis,^10,11^ mainly in the model plant Arabidopsis, where the meiotic regulation is primarily studied in PMCs which can be found at a quantity of 100-120 per male floral organ (anther, with six anthers present per flower) and which are relatively easily accessible. In contrast, female meiosis remains much less known. In Arabidopsis, female meiosis occurs within ovules, which are deeply embedded in maternal floral organs (carpels) and thus are not readily observable. Moreover, the roughly 700 PMCs per flower bud are matched by only approximately 50 MMCs. This low proportion makes it for instance very laborious to find MMCs in squash preparations typically used for immuno-cytochemistry experiments of plant meiosis. Moreover, very little is known about the dynamics of female meiosis and the few studies that have addressed this question have relied on time course experiments with fixed material precluding the tracing of a single cell and an estimation of the biological variation.^12^

Recent advances in live-cell imaging of meiosis have opened new possibilities for observing cellular processes with great spatial and temporal resolution. This has provided important insights into male meiosis, including the analysis of meiotic progression,^13–15^ chromosome movement,^16,17^ meiotic spindle dynamics,^18–20^ positioning of crossovers,^21^ repositioning of the division plane,^22^ and chromosome axis organization.^23,24^

In this study, we have established a live-cell imaging system to visualize female meiosis in Arabidopsis. Based on the difference between female and male meiosis, a new analysis system had to be developed that relied on the formation of a contig of meiotic reporter lines that covered the entire duration of meiosis. This system revealed landmarks of female meiosis and allowed the quantification of meiotic stages with high temporal resolution. With this, we have obtained a temporal and spatial framework of female meiosis that also provides criteria for the evaluation of meiotic mutants. Here, we have used this system to reanalyze quadruple mutants in cyclin-dependent kinase inhibitors which undergo several mitotic divisions before executing meiosis.^25^ This work highlights that a meiotic division can, surprisingly, be converted to a mitotic division with several meiotic structures already established. On the other hand, we found that meiotic fate was already heritably fixed very early in cells destined to undergo meiosis.

## Results

### Set up of a live-cell imaging system for female meiosis in Arabidopsis

To establish a live-cell imaging system for female meiosis in its developmental context, we devised a protocol in which inflorescences were excised from flowering plants and placed on an agar plate. All flower buds were removed except for a single young bud at the meiotic or premeiotic stage, which was left attached to a segment of the inflorescence stem (Figure 1A, B). From this bud, sepals, petals, and stamens were carefully cut off to expose the central pistil. Subsequently, one valve of the ovary was dissected to reveal the ovule primordia containing the MMCs, which were oriented with the MMCs facing toward the objective of the microscope (Figure 1E). The stem of the opened ovary was then embedded in half-strength Murashige and Skoog (MS) medium containing 0.8% agarose and the ovary was further stabilized with a drop of 1% agarose placed on top (Figure 1E). Finally, the petri dish was filled with water. Imaging was performed using a water-dipping objective on an upright confocal laser scanning microscope, focusing on a single primordium (Figure 1F–H).

**Figure 1.**
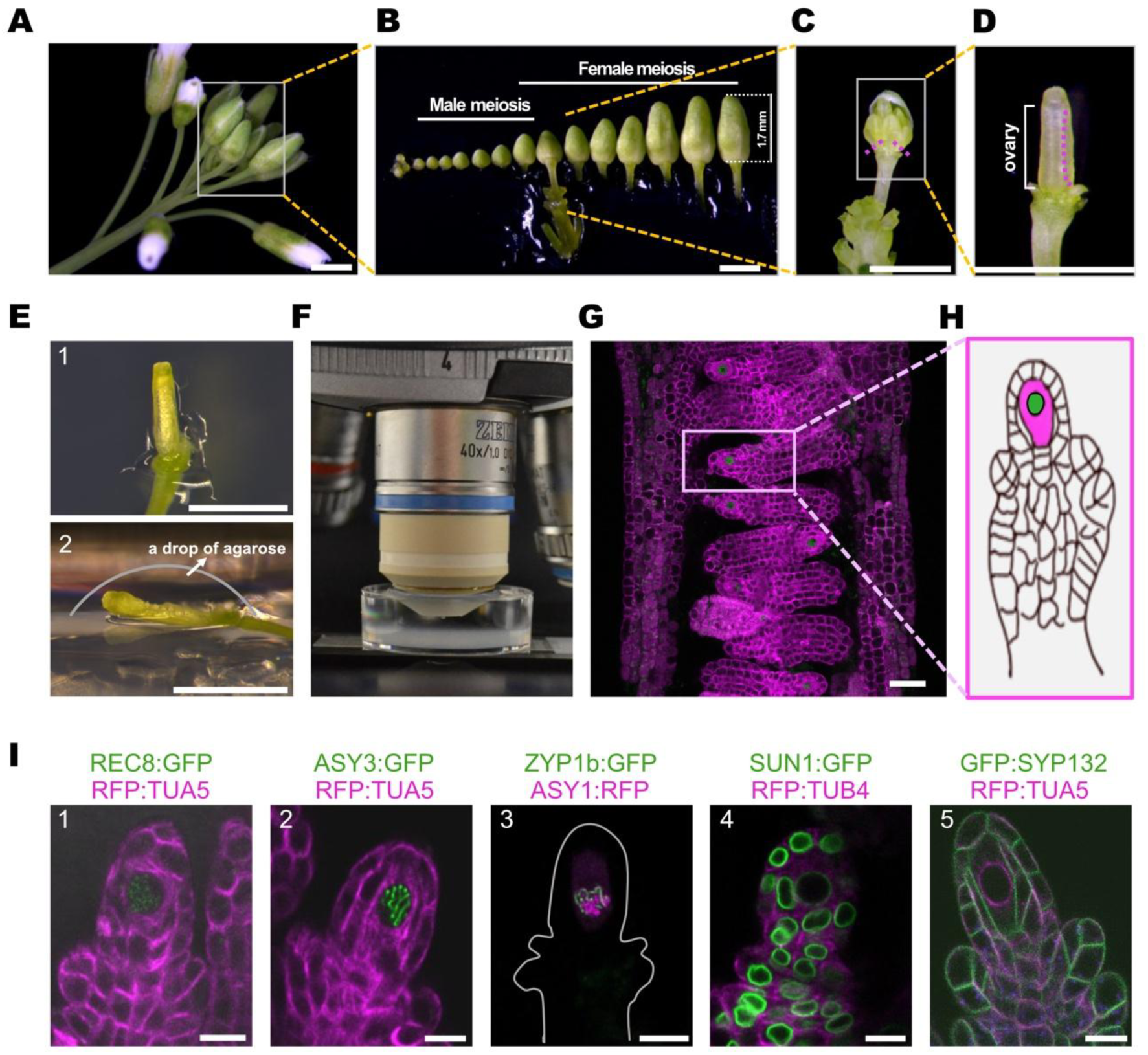
Live-cell imaging setup for female meiocytes. **(A)** Inflorescence of a 5-week-old plant. Scale bar, 1 mm. **(B)** Flower buds at the meiotic stage. Scale bar, 1 mm. **(C)** Removal (magenta dashed line) of sepals, petals, and stamens from a flower bud exposes the ovary. Scale bar, 1 mm. **(D)** Removal (magenta dashed line) of one ovary valve reveals the ovule primordium containing the megaspore mother cells (MMCs). **(E)** Opened the ovary with the stem embedded in the medium. (1) Top view. (2) Lateral view. The opened ovary will be stabilized with a drop of 1% agarose (border of the not yet applied drop is indicated by a grey line). Scale bar: 1 mm. **(F)** The specimen is submerged in water and positioned for imaging with a water-dipping lens. **(G)** Confocal laser scanning micrograph showing several ovule primordia with MMCs highlighted by a meiosis-specific reporter line. Scale bar, 25 μm. **(H)** Scheme of an ovule primordium with the distal MMC highlighted in magenta and its nucleus in green. **(I)** Representative images of meiotic reporter lines used in this study: (1) REC8 (green) and TUA5 (magenta); (2) ASY3 (green) and TUA5 (magenta); (3) ZYP1b (green) and ASY1 (magenta); (4) SUN1 (green) and TUB4 (magenta); (5) SYP132 (green) and TUA (magenta). Scale bar, 10 μm.

Since female and male meiosis are not initiated at the same time during Arabidopsis flower development,^12^ previous estimates on the floral stage when male meiosis takes place could not be used as a proxy for the time point of female meiosis.^13^ We therefore first established a rough developmental time course in which we correlated the length of the ovary with the time point when meiosis occurs revealing that under our growth conditions, female meiosis takes place when the flowerbud is between 0.8 and 1.7 mm long harboring an ovary between 0.5 and 1 mm in a 5-week-old plant (Figure 1B).

Next, we determined that ovules containing MMCs can be kept alive for more than 60 hours with this setup. Since we observed that the same meiotic stages lasted for approximately the same duration irrespective of at what stage a movie was started (Table S1), we conclude that our setup does not, at least not markedly, interfere with meiotic progression, paving the road for a detailed live-cell imaging approach.

### Microtubule dynamics during female meiosis

Cellular changes during meiosis typically occur as a continuum, presenting a challenge for the analysis of live-cell imaging data that relies on quantifying these changes and/or translating gradual, quantitative differences into qualitative criteria to define cytological stages. In previous live-cell imaging studies of Arabidopsis male meiocytes, we observed that the microtubule cytoskeleton is highly dynamic and adopts distinct configurations that can serve as reliable landmarks for staging meiosis.^13^ On the male side, we previously identified 11 discrete microtubule states, such as the formation of a microtubule crescent on one side of the nucleus (the "half moon" stage) and the complete enclosure of the nucleus by microtubules (the "full moon" stage). Thus, even the sole use of a microtubule reporter line enabled a fine-grained dissection of male meiotic progression.^13^

Therefore, we explored the microtubule cytoskeleton during female meiosis with tubulin reporters in which TagRFP is fused to TUB4 or TUA5. However, the microtubule organization during prophase I provided to our surprise only a few criteria to dissect meiosis. At the onset of meiosis, microtubules were homogeneously distributed in the cytoplasm (Figure S1A, Movie S1). In early prophase I, distinct microtubule bundles extend from the spindle poles possibly to anchor the nucleus in a central position (Figure S1A, Movie S1). As meiosis progresses to late prophase I, microtubules accumulate around the nuclear membrane and form a perinuclear ring at diplotene (Figure S1A, Movie S2), resembling the “full-moon” structure previously described in different plant species including Arabidopsis,^13^ potato,^26^ wheat and tomato.^27^

Following nuclear envelope breakdown, the perinuclear ring collapses, and microtubules straighten to invade the former nuclear space. These microtubules reorganize into bipolar spindle fibers, which align the chromosomes at the metaphase plate (Figure S1A, Movie S2). The spindle fibers converge at the poles and extend in parallel along the division axis. During anaphase I, kinetochore-attached microtubules shorten, drawing homologous chromosomes toward opposite poles. Upon reaching the poles at telophase I, the spindle disassembles and nuclear envelopes reform around the segregated chromosomes. Concurrently, the phragmoplast forms between the daughter nuclei, with parallel microtubules guiding cell plate formation. As division progresses, the phragmoplast expands centrifugally until the cell plate fuses with the parental plasma membrane (Figure S1A, Movie S2).

During interkinesis, curved microtubules accumulate around the two nuclear membranes, forming two perinuclear rings (Figure S1B, Movie S2). Upon nuclear envelope breakdown, these perinuclear rings collapse simultaneously, and microtubules straighten to invade the former nucleus space, assembling two spindle apparatuses oriented in the same direction as in metaphase I (Figure S1B, Movie S2). During the second meiotic division, microtubules in each cell behave similarly as in metaphase to telophase of meiosis I (Figure S1B, Movie S2). At anaphase II, sister chromatids separate towards opposite poles, and spindle structures subsequently disassemble. Nuclear envelopes reform around the four sets of segregated chromosomes, while two phragmoplasts are deposited from the center simultaneously between each newly reformed nucleus (Figure S1B, Movie S2). As cytokinesis progresses, these two phragmoplasts expand laterally along the division plane until they fuse with the parental plasma membrane, like the cytokinetic pattern of meiosis I.

We conclude that a microtubule reporter can be used to define distinct stages from metaphase I onwards, primarily due to chromosome separation and cell division structures such as the spindle and the phragmoplast (Figure S1A, B, Movie S2). However, this reporter only provides limited temporal resolution during prophase I. The reason for this is that the MMC, while growing, maintains a club-shaped appearance over the entire time course of meiosis (Figure S1A, see below). This is in contrast to the male meiocyte, which changes during prophase from round via oval to trapezoidal and finally tetrahedral, a change that can be precisely followed by observing MTs. In addition, characteristic male microtubule structures, foremost a “half moon” arrangement flanking the nucleus, were absent.

### Analysis of further reporter lines to dissect female prophase I

Recognizing the limited resolution of a microtubule reporter, we sought an alternative system to monitor meiotic progression through prophase I. In the past, we and others have generated several fluorescently tagged functional genomic reporter lines for many meiotic regulators,^28^ including REC8,^13^ ASY1,^14,24,29^ ASY3,^24,29^ and ZYP1.^24^ In addition, we and others have also constructed many other reporter lines that allow monitoring additional cellular features, such as SUN1 for the nuclear membrane^30,31^ and SYP132 for the plasma membrane.^20,32^ Therefore, we explored the possibility of monitoring female meiosis with these reporters.

REC8 is a meiosis-specific subunit of cohesin, which holds the sister chromatids together and extrudes chromatin loops.^33^ REC8 is also a crucial component of the chromosome axis and plays a key role in recruiting other axis proteins, such as ASY3.^34^ Monitoring REC8 can therefore be used to track chromosomes and monitor their organization. At the same time, the nucleolus can be seen as an area in the nucleus without a REC8 label.

Similar to male meiosis,^13^ we found that REC8 accumulates very early in female meiocytes. REC8:GFP appears as a diffused dotty signal within the nucleoplasm in early prophase I (Figure 2A, Movie S1). As REC8 loading continues and chromosomes start to condense, the GFP signal intensifies, forming thin thread-like structures (Figure 2B, Movie S3). These threads thicken from zygotene to pachytene, indicating synapsis and further chromosome condensation. Concomitant with synapsis, REC8-marked chromosomes move rapidly in the nucleus (Movie S3).

**Figure 2.**
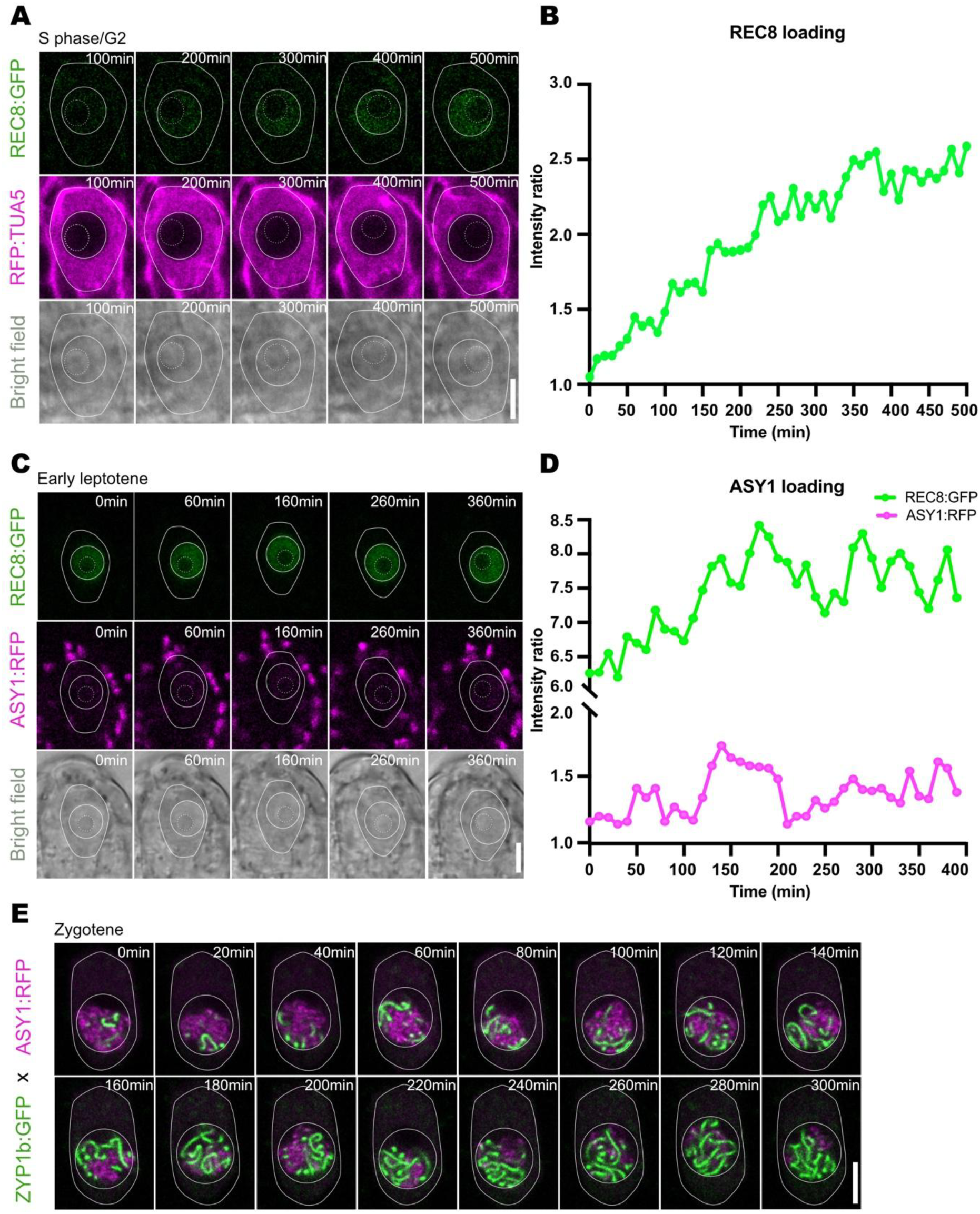
Visualization of chromatin dynamics in female meiocytes using reporter lines. **(A)** Loading of REC8:GFP onto chromatin in female meiocytes. Scale bar, 5 μm. **(B)** Fluorescence intensity ratio of nuclear to cytoplasmic REC8:GFP during REC8 loading in (A). **(C)** Simultaneous analysis of ASY1:RFP and REC8:RFP loading in female meiocytes. Scale bar, 5 μm. **(D)** Fluorescence intensity ratios of nuclear to cytoplasmic REC8:GFP and ASY1:RFP during ASY1 loading in (C). **(E)** Synaptonemal complex assembly visualized by the loading of ZYP1b:GFP and the removal of ASY1:RFP during synapsis. Scale bar, 5 μm.

As seen by REC8:GFP, chromosomes further condense as meiosis progresses and the REC:GFP signal becomes dim, likely due to the removal of REC8 by WAPL, marking the disassembly of the synaptonemal complex and homolog desynapsis.^23^

At diakinesis, chromosomes are condensed into short, rod-like structures, the nucleolus disintegrates. After the nuclear envelope breakdown, we could detect REC8:GFP signals aligning in the middle of the spindle microtubules during metaphase I (Movie S2). However, we could not visualize chromosome behavior after homolog segregation at anaphase I using the REC8 reporter presumably because only a few cohesion molecules remain at the centromeres, providing not sufficient fluorescence to be imaged.

Following the chromatin dynamics with REC8:GFP also gives an approximation of where the nucleus is located within the cell. In contrast to male meiocytes, we found that the nucleus in the MMC resides in a central position, although some cell-to-cell variation could be observed. At the same time, the position of the nucleolus within the nucleus can be deduced by the absence of REC8:GFP fluorescence (Figure 2A). When REC8 accumulation starts, the nucleolus is about half the size of the nucleus and gradually migrates off-center. Then, it continues to move toward the periphery, touching the nuclear envelope by late leptotene and rotating together with the envelope throughout the rest of prophase (Movie S3).

ASY1 is a chromosome axis-associated HORMA-domain protein, which plays a crucial role in Arabidopsis in CO positioning, interference and assurance.^35–37^ ASY1:RFP is clearly visible in early prophase before the nucleolus touches the nuclear periphery (Figure 2C). ASY1 and REC8 loading appear to overlap with REC8 likely starting earlier than ASY1 to be recruited (Figure 2D). However, due to the weak fluorescence of ASY1:RFP, it was difficult to unambiguously determine the exact time point of the beginning of ASY1 loading.

ASY1 presence on chromosomes is temporally restricted and with the installation of the synaptonemal complex, which establishes a proteinaceous connection between the homologous chromosomes, it is removed from the axis ^36,37^. The formation of the synaptonemal complex can be visualized by the accumulation of the transverse filament protein ZYP1, which polymerizes between the paired chromosome axes. Therefore, we also combined the ASY1 reporter (PROASY1:ASY1:TagRFP) with a ZYP1 reporter (PROZYP1b:ZYP1b:GFP). This allowed us to visualize the onset of synaptonemal complex formation, marked by the accumulation of ZYP1b:GFP in short stretches that progressively elongate into long filaments, and its full installation indicated by the absence of ASY1:RFP (Figure 2E, Movie S4).

Chromosome pairing, recombination and synapsis have been found to be associated with fast chromosome movements.^16,17^ Chromosome movements are driven by external forces, generated by microtubules and their associated proteins as well as the actin-myosin cytoskeleton, which connects to nuclear envelope proteins. With the help of the LINKER OF THE NUCLEOSKELETON AND CYTOSKELETON (LINC) complex, the external force is transmitted through the nuclear envelope to drive chromosome movements within the nucleus. SUN1, a Sad1/UNC-84 (SUN) domain protein, which localizes to the inner nuclear membrane, is a central component of the LINC complex and tethers telomeres to the nuclear envelope during meiosis.^17,31,38^ Thus, the SUN1 distribution pattern is a proxy for the attachment of telomeres to the nuclear envelope driving chromosome movement and bouquet formation.

The published SUN1 genomic reporter, PROSUN1:SUN1:GFP,^30,31^ was combined with PRORPS5A:TagRFP:TUA5. SUN1:GFP localizes uniformly in the nuclear envelope, forming a wrinkled ring in premeiotic cells when the nucleolus is centrally located, then gradually tenses as meiocytes progress into leptotene and the nucleolus moves off center (Figure 3A). From late leptotene, SUN1:GFP displays a heterogeneous distribution, with several intensely labeled domains interspersed within an otherwise uniform SUN1:GFP signal (Figure 3B, white arrows). Notably, approximately one-third of the nuclear envelope shows a markedly reduced SUN1:GFP signal, where only a faint amount remains detectable (Figure 3B).

**Figure 3.**
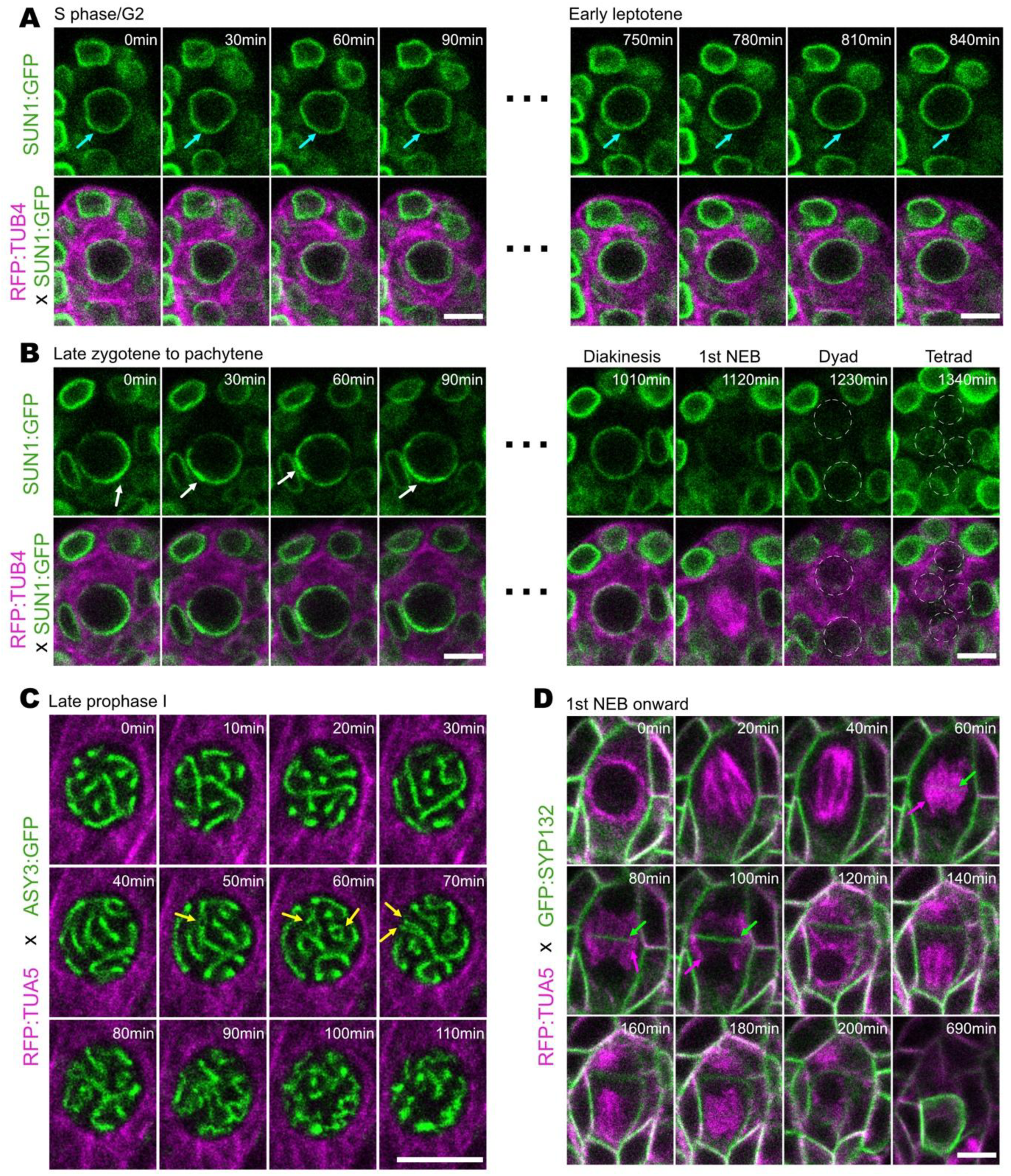
Visualization of nuclear envelope organization, chromosome dynamics, and cell wall formation in female meiocytes. **(A)** Live-cell imaging of PROSUN1:SUN1:GFP × PRORPS5A:TagRFP:TUA5 in a female meiocyte during early prophase. The nuclear envelope (blue arrow) is wrinkled in S phase/G2, reflecting the appearance of the nuclear envelope of cells surrounding the MMC, and becomes smoother as meiosis progresses into early leptotene while the nuclear envelope of the surrounding cells maintains its wrinkled appearance. Scale bar, 5 μm. **(B)** Live-cell imaging of PROSUN1:SUN1:GFP × PRORPS5A:TagRFP:TUA5 in a female meiocyte during late prophase. SUN1:GFP signal is depleted from one-third of the nuclear envelope and intensifies (white arrow) in distinct domains during zygotene and pachytene. In diplotene, SUN1 is uniformly distributed in the nuclear envelope. Scale bar, 5 μm. **(C)** Live-cell imaging of PROASY3:ASY3:GFP × PRORPS5A:TagRFP:TUA5 in a female meiocyte at diplotene. During the diplotene quiescence period, rapid chromosome movement ceases and chromosomes become immobile as the synaptonemal complex disassembles. Yellow arrow: split homologous chromosomes. Scale bar, 5 μm. **(D)** Live-cell imaging of PROSYP132:GFP:SYP132 × PRORPS5A:TagRFP:TUA5 in a female meiocyte during division. Phragmoplast formation (magenta arrow) and inside-to-outside cellplate expansion (green arrow) are observed during successive cytokinesis. SYP132 signal intensifies in only one meiotic product, the functional megaspore, after tetrad formation. Scale bar, 5 μm.

By late pachytene, SUN1 distribution became uniform around the entire nucleus again and remained unchanged until nuclear envelope breakdown, after which SUN1:GFP signals were undetectable (Figure 3B). The nuclear envelope was absent during metaphase I and anaphase I, but reformed around decondensed chromosomes at telophase I, with SUN1:GFP uniformly reappearing at the nuclear periphery of each daughter cell (Figure 3B).

Since REC8:GFP signal becomes diffuse in late prophase (see above), we also followed the axis protein ASY3 to monitor chromatin movement using the previously published reporter line PROASY3:ASY3:TagRFP.^23,24^ This reporter allowed especially well the visualization of the rapid movements of chromosomes prior to and during synapsis (Movie S5). Remarkably, when the synaptonemal complex starts to be disassembled, but before nuclear envelope breakdown, the rapid chromosome movement comes to a halt that persists for nearly two hours. The thick thread-like appearance of chromosomes marked by ASY3:GFP concomitantly split into two thinner, yet juxtaposed lines (Figure 3C, yellow arrows), indicating the disassembly of the synaptonemal complex. In the following, chromosomes further condense into rod-like structures in diakinesis (Figure 3C, Movie S6).

SYP132 is a SNARE protein (SOLUBLE N-ETHYLMALEIMIDE-SENSITIVE FACTOR ATTACHMENT PROTEIN RECEPTOR), needed for vesicle fusion and localized at the plasma membrane in all plant tissues.^20,32^ Using a SYP132 reporter line enables the precise observation of cytokinesis as it marks the cell plate that starts to form at the center of the cell between the newly formed nuclei, growing outwards until it fuses with the parental plasma membranes producing a temporary dyad after the first meiotic division (Figure 3D, Movie S7). Using GFP-tagged SYP132 in combination with RFP-tagged TUA5, we clearly visualize the formation of two daughter cells after cytokinesis, oriented along the nucellus-chalaza direction (Figure 3D, Movie S7).

Ultimately, four haploid daughter cells are generated in a linear tetrad within the ovule (Movie S7). Approximately three to four hours post-division, three of these meiotic products undergo programmed cell death, while the one positioned at the chalaza end survives, marked by an enrichment of SYP132 fluorescence (Figure 3D). The surviving megaspore subsequently enlarges, occupying also the space of the degenerated cells.

### Defining landmarks of female meiosis by a reporter contig system

Analyzing the above-described reporters allowed us to identify distinct events during female meiosis. We refer to these events as landmarks. In total, we were able to define 15 landmarks (**L1** to **L15**, Figure 4). Importantly, a defined order of these landmarks could be established as the features and durations covered by our reporters overlap building a contig that completely covers female meiosis. A crucial aspect of this analysis was that the overlapping movies can be cross-anchored for three reasons: First, different reporters sometimes mark the same feature, e.g., REC8 and ASY1 can both be used to highlight chromosomes until synapsis. Second, most of our reporter lines express a tubulin reporter next to a reporter for another meiotic gene, e.g., *PROREC8:REC8:GFP and PRORPS5ATagRFP:TUA5.* Thus, even though tubulin has only a limited resolution for meiosis I, it still highlights a few features that can be used to connect different movies, especially a the *full moon* structure (Figure 1H), Finally, using the bright field channel the nucleolus position and the presence/absence of the nuclear envelope can usually be determined. This resulted in the order of landmarks depicted in Figure 4A for prophase I and Figure 4B for the second meiotic division.

**Figure 4.**
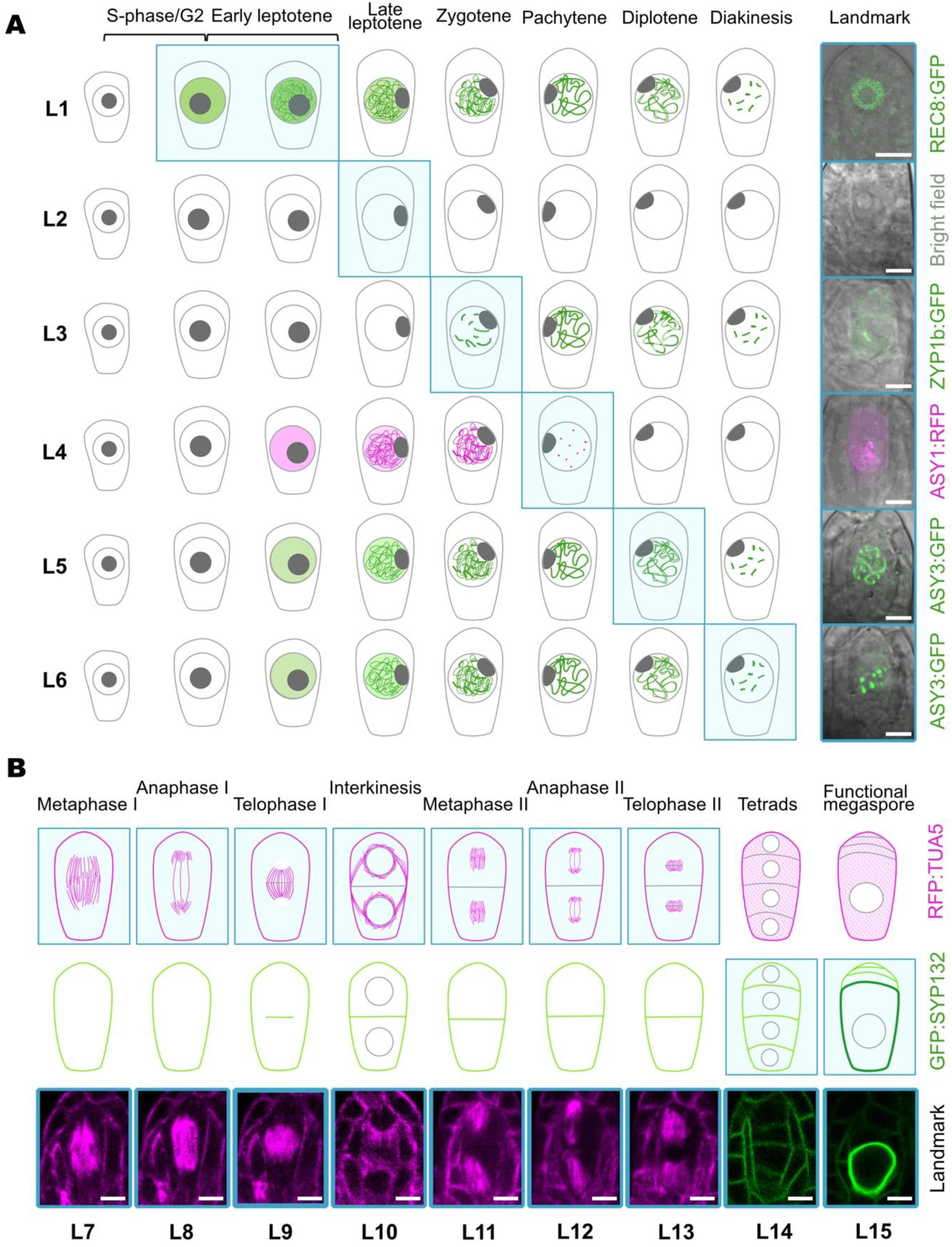
Schematic overview of the 15 meiotic landmarks (L1–L15). **(A)** Landmarks L1–L6 (indicated in blue boxes) are defined by chromosome behaviors and cytological features. The final column displays representative microscopy images of meiocytes at each stage. L1 is defined by the loading of REC8, corresponding to late G2 and early leptotene. L2 marks the onset of late leptotene, identified by the nucleolus touching the nuclear envelope. L3 indicates entry into zygotene, marked by the loading of ZYP1b onto chromosomes. L4 represents the extensive removal of ASY1 at early pachytene. L5 marks the onset of diplotene, coinciding with the cessation of chromosome movement and synaptonemal complex disassembly. L6 indicates the beginning of diakinesis, when chromosomes condense into dot-like structures. Scale bar, 5 μm. **(B)** Landmarks L7–L15 (indicated in blue boxes) are characterized by specific microtubule configurations and cell wall formation during meiosis and cytokinesis. The bottom row provides microscopy images corresponding to each landmark. Scale bar, 5 μm.

Prior to meiosis, the destined female meiocyte in Arabidopsis and also other plant species expands making it usually the largest cell at the distal (chalazal) end of an ovule primordium. Likely associated with cell growth, the meiocyte shows gradually thickening cortical microtubules (Figure S1). At this stage, the nucleus is positioned centrally and the nucleolus is present in the center of the nucleus (Figure S1). The first landmark is the beginning of the loading of REC8 onto chromosomes and the beginning of chromosome condensation so that the chromosomes become visible as thin, thread-like structures (**L1**). Based on work in yeast and animals, the loading of REC8 likely takes place in the *premeiotic S-phase* or *early G2- phase*.^39,40^ The second landmark (**L2**) is a repositioned nucleolus, which moves from a center position to the side of the nucleus. This is made possible through the release of chromosomes from the nuclear envelope allowing them to pair;^41^ the landmark defines *late leptotene*. *Zygotene* is demarcated by the beginning of the installation of the synaptonemal complex leading to synapsis, which can be visualized by the loading of ZYP1 along the chromosome axes (**L3**). As synapsis progresses, the chromosome axis-associated protein ASY1 starts to be widely removed from the chromosome axes and chromosomes further condense (**L4**), marking *pachytene* (Figure 4B). Chromosomes exhibit fast movement from late leptotene onwards, possibly facilitating homolog recognition, pairing and recombination (Movie S3). Interestingly, this movement becomes more rapid in *zygotene* and *pachytene*. However, chromosome movement suddenly slows down to an almost complete arrest likely through the re-anchoring of chromosomes to the inner nuclear membrane;^17,41^ at the same time, homologous chromosomes begin to desynapse defining **L5** and *diplotene*. Chromosomes condense further in diakinesis, and the linear threads of the chromosome axis protein ASY3 compacted into short rods (**L6**), appearing as highly condensed bivalents. Then, the nuclear envelope breaks down marking the end of prophase I.

At *metaphase I*, the microtubule spindle apparatus appears in the middle of the cell (**L7**), reflecting the highly condensed chromosomes aligned along the metaphase plate. The transition to *anaphase I* is marked by thinner and elongated spindle shape, with accumulation of microtubules close to two spindle poles, indicating the shortening of spindle fibers separating the homologous chromosomes toward opposite poles (**L8**). As the chromosomes reach the poles, the spindle disassembles, and the nuclear envelopes begin to reform around the segregated chromosomes. The phragmoplast between the two daughter nuclei starts to form in *telophase I* highlighted by microtubules aligning in the middle of the cell (**L9**). As the first division proceeds, the phragmoplast dissolves from the middle marking the margin of the cell plate until the newly formed plasma membranes separate the first two meiotic products. In parallel, two perinuclear ring structures formed by microtubules appear (**L10**). Shortly after, meiosis II commences and the nuclear envelope of the two daughter nuclei breaks down. The two spindles of the second meiotic division are formed by microtubules (**L11**) marking *metaphase II*. **L12** represents *anaphase II* and is visualized by two thinner spindles aligned at the same direction, with microtubules accumulated at the spindle poles, which pull the homologous chromosomes to the opposite poles within each daughter cell simultaneously. Two phragmoplasts form in the middle of each daughter cell (**L13**) indicating *telophase II*. At *telophase II*, four nuclear envelopes form around the segregated chromatids. Then *cytokinesis II* results in the formation of a linear tetrad of four haploid daughter cells highlighted by GFP:SYP132 (**L14**). Finally, only the meiotic product that is closest to the basal (micropylar) end of the ovule primordium survives and starts to strongly accumulate SYP132 at the plasma membrane (**L15**), gradually filling the space of the former meiocyte and representing the differentiation of the functional female megaspore, which will then undergo three mitotic divisions in Arabidopsis to form the embryo sac.

### Time course of female meiosis

With the landmark-based classification of different meiotic stages, we could then estimate the time frame of female meiosis in Arabidopsis (Figure 5A). To this end, 105 movies were analysed that captured the transition from one landmark to the next.

**Figure 5.**
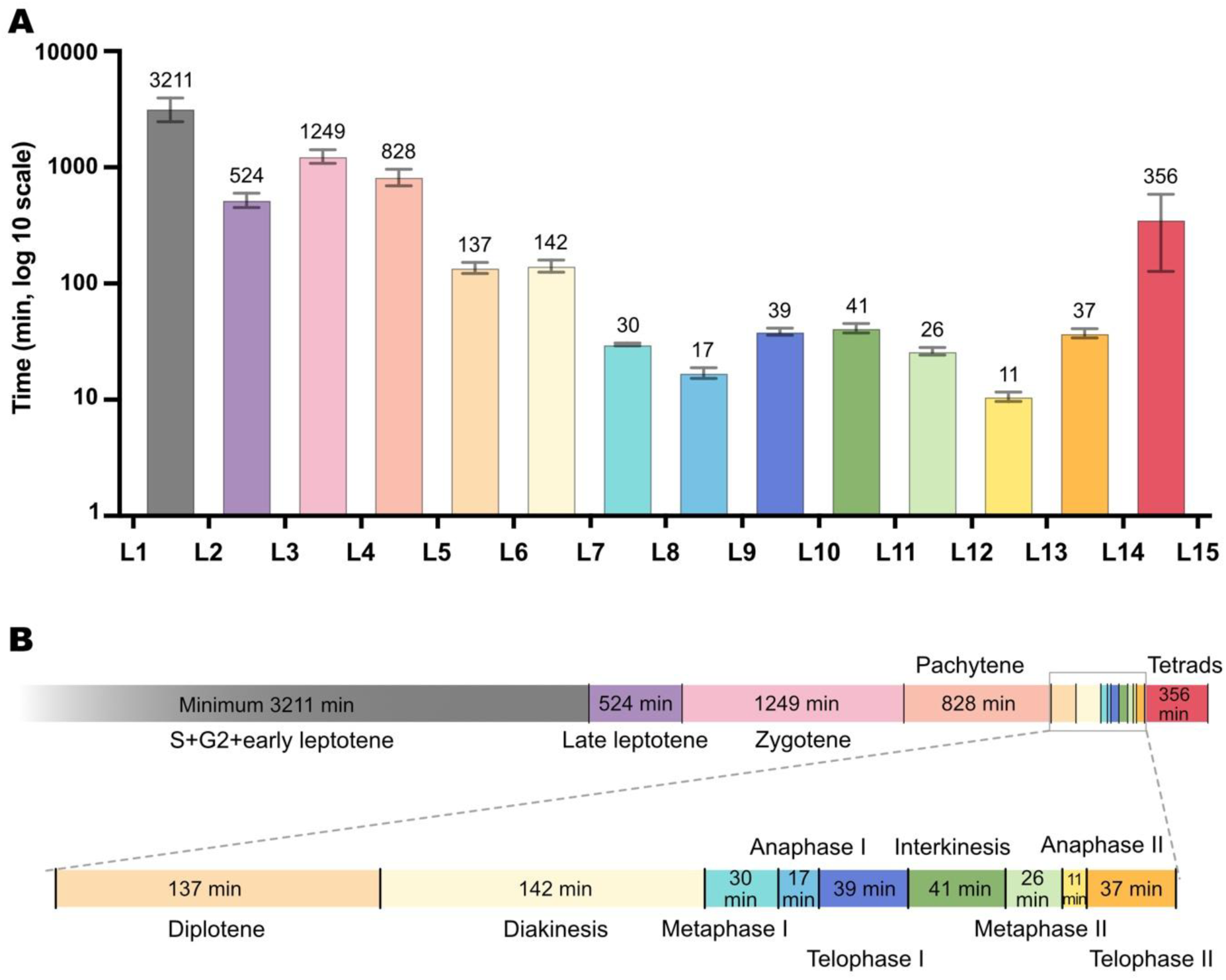
Time course of female meiosis. **(A)** Duration of each stage (in minutes) between consecutive landmarks in wild-type (WT) plants. Column represents the duration on a log10 scale. The median stage duration is indicated at the top of each column, and error bars represent the 95% confidence interval. Bars at the bottom represent the actual duration of each stage. Grey: S+G2+early leptotene; Purple: late leptotene; Pink: zygotene; Coral: pachytene; Light orange: diplotene; Light yellow: diakinesis; Aqua blue: metaphase I; Sky blue: anaphase I; Navy blue: telophase I; Green: interkinesis; Light green: metaphase II; Bright yellow: anaphase II; Orange: Telophase II; Red: tetrads. (**B**) Linear representation of the durations shown in (A). Only the median times are depicted in each bar. The color code is the same as in (A).

As MMCs from one carpel cannot be regarded as statistically independent measurements, the statistical analysis had to account for this clustered structure. Since the observed MMCs sometimes move out of the focal plane, the exact start and/or end of each state could not always be observed. Thus, the data were considered censored. Following the statistical approach previously described in male meiosis,^15^ meiotic phase-specific parametric models for censored time-to-event data were used as the duration of phases is independent of each other. For detailed description of the models, please see the subsection “Statistical methods” in the section “Materials and methods”.

Below, we provide the average lengths between the landmarks of female meiosis and to which “classical” meiotic phase they roughly correspond. The minimal duration of the phase between the premeiotic G1 and S-phase and the first landmark (L1) was at least 590 minutes, nearly 10 hours. This estimate is based on the increase in cell size and the rearrangement of the microtubule cytoskeleton as seen very early during female germline development ^43^. As meiosis progressed, the phase **between L1 and L2** (representing early leptotene) was 3211 minutes (95% CI 2466-3956 min,n=31 MMCs). **L2 to L3** (late leptotene) lasted approximately 524 minutes (95% CI 450-599 min,n = 34). Zygotene duration (**L3 to L4**) was estimated to be 1249 minutes (95% CI 1081-1417min, n = 30). The pachytene stage (**L4 to L5**) lasted 828 minutes (95% CI 692-964 min, n = 40). Diplotene (**L5 to L6**) lasted around 137 minutes (95% CI 122-152 min, n=37). The duration of diakinesis (**L6 to L7**) was about 142 minutes (95% CI 125-159 min, n=36). With the nuclear envelope breaking down, metaphase I (**L7 to L8**) lasted for 30 minutes (95% CI 29-31 min, n = 34). The phase **between L8 and L9** (anaphase I) was 17 minutes (95% CI 15-19 min, n=34). **L9 to L10** (telophase I) lasted about 39 minutes (95% CI 36-41 min, n=34). The phase **between L10 and L11** (interkinesis I) lasted around 41 minutes (95% CI 37-45 min, n=34). Then meiosis II starts. The duration of meiosis II division is shorter than that of meiosis I. The metaphase II duration (**L11 to L12**) was estimated around 26 minutes (95% CI 24-28 min, n=34). The anaphase II (**L12 to L13**) lasted about 11 minutes (95% CI 10-12 min, n=34). The telophase II (**L13 to L14**) was followed around 37 minutes (95% CI 34-41 min, n=34). Then the cytokinesis happens at the same time for the two daughter cells. The linear tetrads (**L14 to L15**) lasted 356 minutes (95% CI 127-585 min, n=33), where the survival female microspore is enriched with brighter SYP132 signaling (Figure 4B-C).

Adding these durations, we estimated that the total median duration of female meiosis from early leptotene (L1) to the appearance of the functional megaspore (L15) is around 112 hours, more than four and a half days. Prophase I (L1 - L6) was found to be particularly long, almost 103 hours, i.e., representing more than 90% of the total duration. Division stages of meiosis I and meiosis II are much shorter compared to prophase I. The duration of metaphase is nearly the same in meiosis I and meiosis II (Figure 5B).

### Nuclear size of the MMC increases during prophase

The nuclear size of male meiocytes has been found to increase during prophase I.^17^ Thus, we wanted to know whether an increase could also be observed on the female side. We determined that the diameter of the MMC nucleus at leptotene is 6.8 ± 0.7 μm on average, 8.2 ± 0.9 μm at zygotene/pachytene and 8.5 ± 0.6 μm at diakinesis (Mean ± SD, n=28, 22 and 27 respectively). Assuming a spherical shape the volume increases from an average of 164.6 μm^3^ via 288.5 μm^3^ to 321.6 μm^3^ representing a total increase of 95%.

The nucleus of the MMC is much larger than the nuclei of the surrounding epidermal and subepidermal cells in an ovule primoridium.^42^ However, when we measured the size of the nucleus of the male meiocyte, we found that its diameter at leptotene is 7.2 ± 0.8 μm, 8.3 ± 0.9 μm at zygotene/pachytene and 8.9 ± 0.7 μm at diakinesis (Mean ± SD, n=27, 51 and 21 respectively), which matches the measurement from Cromer et al. (2024). With that, the nucleus of the MMC is comparable in size to the nucleus of the male meiocyte in leptotene, zygotene/pachytene and diakinesis, with no statistically significant difference observed (p > 0.05).

### Determination of meiotic commitment

Having established a temporally resolved cytological framework for female meiosis, we next focused on a central question of female reproductive development, i.e., when is an MMC committed to undergo meiosis. To address this question, we have analyzed the quadruple mutant *krp3 krp4 krp6 krp7* in which a family of CDK inhibitors (KIP27-RELATED PROTEINS, KRPs) is defective. In these mutants, the designated MMC undergoes several mitotic divisions prior to meiosis, resulting in the formation of supernumerary meiocytes and subsequently in multiple gametophytes within a single ovule.^25^

Given the crucial role of REC8 in the regulation of cohesion during meiosis and, consequently, in chromosome segregation, we hypothesized that the replacement of RAD21 by the meiosis-specific REC8 kleisin variant could be pivotal for meiotic progression. To test this, we co- introduced PROREC8:REC8:GFP and PRORPS5A:TagRFP:TUA5 into the *krp3 krp4 krp6 krp7* mutant background. The resulting transgenic plants were subjected to live-cell imaging. In total, 71 ovules from 64 independent plants were analyzed. Five movies showed a mitotic division of a presumptive MMC prior to REC8 loading as judged by the size of the cell and the nucleus (Movie S8). However, we did not see REC8:GFP accumulating afterwards making it difficult to judge the fate of these cells. Two other presumptive MMCs showed neither detectable REC8 loading nor any sign of cell division. In 64 movies, REC8:GFP expression was captured. Among these, 33 showed no evidence of cell division, 14 showed meiocytes progressing through meiosis while 17 movies depicted meiocytes undergoing a mitotic division (Figure 6A). The daughter cells resulting from the mitotic divisions exhibited stronger REC8:GFP signals than the mother cell (Movie S9), indicating that, at least in part, meiotic protein synthesis was maintained and REC8:GFP was not degraded or replaced. The continued production of REC8:GFP, indicated that these cells still have MMC fate. These findings suggest that, although meiotic commitment appears to be established early, i.e, before REC8 accumulation, a meiotic division can be replaced by a mitotic division after REC8 loading. Conversely, a mitotic division can be executed in the presence of this kleisin subunit being loaded in place of RAD21.

**Figure 6.**
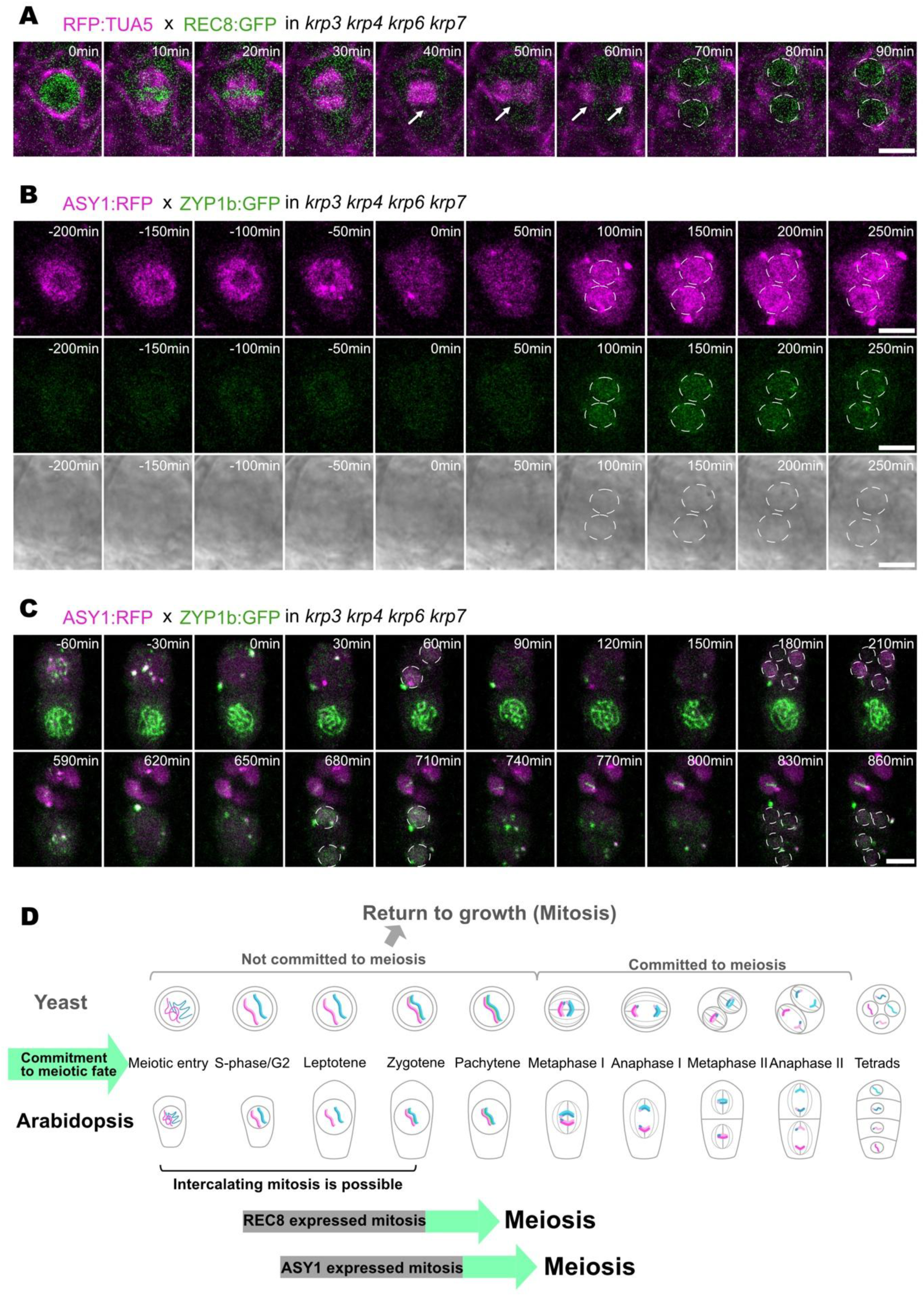
Analysis of meiotic commitment in *krp3 krp4 krp6 krp7* mutants. **(A)** An MMC of a *krp3 krp4 krp6 krp7* mutant plant with REC8 loaded undergoes one complete division resulting in two separated cells as visualized by the formation of a phragmoplast (white arrow). Notably, REC8 re-accumulated onto chromosomes in both daughter nuclei (highlighted in white dashed circles). Tubulin marked by TUA5 in magenta. Nuclear envelope breakdown marked as 0 min. Scale bar: 5 μm. **(B)** Meiocyte with ASY1:RFP loaded onto chromosomes undergoes one division resulting in two products (highlighted in white dashed circles) expressing both ASY1:RFP (magenta) and ZYP1:GFP (green) in *krp3 krp4 krp6 krp7*. Nuclear envelope breakdown marked as 0 min. Scale bar: 5 μm. **(C)** Two MMCs of a *krp3 krp4 krp6 krp7* mutant plant expressing ASY1:RFP (magenta) and ZYP1:GFP (green) go through two rounds of division in resulting in four products from each meiocyte. Upper meiocyte started first division at 0 min, two products (white dashed circles) expressing ASY1:RFP (magenta), and the second NEB at 90 min leading to four products (white dashed circles) with ASY:RFP (magenta). The lower meiocyte followed the meiotic division, with first NEB at 620 min and second NEB at 740 min. Division products highlighted with white dashed circles. Scale bar: 5 μm. **(D)** Model of meiosis progression in yeast and *krp3 krp4 krp6 krp7*. In yeast, cells can exit meiosis return to mitosis before the commitment in prometaphase I. In Arabidopsis, after the cell commits to a meiotic fate, intercalating mitosis with meiotic structures is possible until zygotene, but it cannot exist meiosis.

Next, we assessed whether MMCs can still mitotically divide when later meiotic structures are established, i.e., the chromosome axis and the synaptonemal complex. To analyze this, we introduced PROASY1:ASY1:TagRFP along with PROZYP1b:ZYP1b:GFP into the *krp* quadrupel mutants. Among 31 movies analyzed, 28 showed detectable ASY1:RFP expression at imaging onset. Two movies showed ASY1-positive meiocytes at early leptotene undergoing a mitosis-like division that produced two daughter cells retaining ASY1 expression; in one case, ZYP1 signals were also detected in the division products (Figure 6B). In an additional movie, two MMCs were present at the imaging onset: the lower MMC gradually acquired ASY1 loading, while the upper one did not (Movie S10). Both MMCs subsequently underwent a mitosis-like division, with the lower-derived products showing stronger ASY1 signal than the mother cell while the products from the upper MMC with no detectable ASY1 (Movie S10). In 16 movies, both ASY1 and ZYP1 were detected, and at least one meiocyte proceeding through meiosis (Figure 6C). In 11 movies, the ASY1 signal diminished in some or all meiocytes during imaging, including two cases where the loss occurred in the mitosis-like division products (Movie S10). Among the three movies lacking ASY1 expression, one movie showed a mitosis-like division generating two daughter cells without detectable ASY1 expression. These findings show that even ASY1 loading and with this an at least partially established chromosome axis is still compatible with mitosis and that a meiotic division is not fixed even after approximately 50 hours into meiosis.

In contrast, in all 16 movies where ZYP1 loading was observed in one or more MMCs in one ovule primordium, ZYP1-positive cells did not undergo a mitotic division and invariably progressed through meiosis until tetrad formation (Figure 6C, Movie S11). Thus, we conclude that a meiotic division is not fixed till at least zygotene. Since we currently cannot judge whether the mitosis-promoting force diminishes over time in *krp3 krp4 krp6 krp7*, we cannot exclude that it might be even possible to induce a mitosis after zygotene, possibly with a reduced probability that would hence require a much larger sample size than ours to be detected. For the same reason, it is currently difficult to determine whether the synaptonemal complex itself represents a determinant of a meiotic division or whether its establishment reflects just the time until when division can be reprogrammed.

Taken together, meiotic fate appears to be heritably established prior to REC8 loading. Mitotic divisions can replace a meiotic division at least until the synaptonemal complex is established. However, the fate of the daughter cells of a presumptive MMC that undergoes mitosis appears to be still meiotic demonstrating on the one hand a great level of developmental flexibility but on the other hand a fixed developmental program.

## Discussion

Female and male meiosis differ in many aspects in most if not all eukaryotic species. However, the insight into female meiosis largely lags behind the knowledge gained on the male side. Understanding female meiosis and comparing it with male meiosis is not only important to get crucial insight into female reproductive development but also to extract general principles of meiotic regulation in one species and between different species. To approach the knowledge gap between female and male meiosis, we have established a live-cell imaging system for MMCs in the flowering plant *Arabidopsis*. This system is based on confocal laser scanning microscopy and a reporter contig-based approach allowing the dissection of female meiosis with great temporal resolution. With our setup, we could keep ovule primorida alive for more than 60 hours with an image taken every ten minutes. In total, we could distinguish 15 distinct meiotic stages, referred to as landmarks. Assigning these landmarks to the classical stages of meiosis allowed us to obtain statistically relevant time intervals for these stages, which led to an estimated total length of at least 112 hours for female meiosis. Our work establishes a cytological framework of female meiosis, spanning from the meiotic G2 phase to the formation of a functional megaspore after telophase II.

### Female versus male meiosis

The cytological framework established for female meiosis allowed us to compare meiotic differences between sexes in Arabidopsis with unprecedented spatio-temporal resolution. For this comparison, we primarily drew on cytological data from our previous analysis of male meiosis, as both studies shared similar growth conditions and imaging methodologies.^13^ While we provide new cytological insights into female-specific features—such as spindle formation and the orientation of division planes leading to a linear arrangement of meiotic products—our main focus is on previously unrecognized aspects of female meiosis.

A first striking difference that became clear in our study is the much longer duration of female versus male meiosis (minimum of 112 hours versus 34.5 hours) (Figure 7A). One reason for this is that premeiotic S-phase, G2-phase and early leptotene appear to be much longer in females than in males (approximately 55h versus 9h). It is currently difficult to shed light on the premeiotic phases due to still lacking reporters. Interestingly, the difference in these early stages closely reflects the difference in the onset of female versus male meiosis in young flower buds,^28^ i.e., approximately 40-50 hours, suggesting that the developmental trigger to initiate meiosis in both species could be synchronous, and possibly be even the same. One possible reason for the longer time to start meiosis in the female versus male could be that MMC specification relies on the selection of one cell from a group of germline-competent cells^43,44^ and that this selection/fixation of cell fate may take additional time.

**Figure 7.**
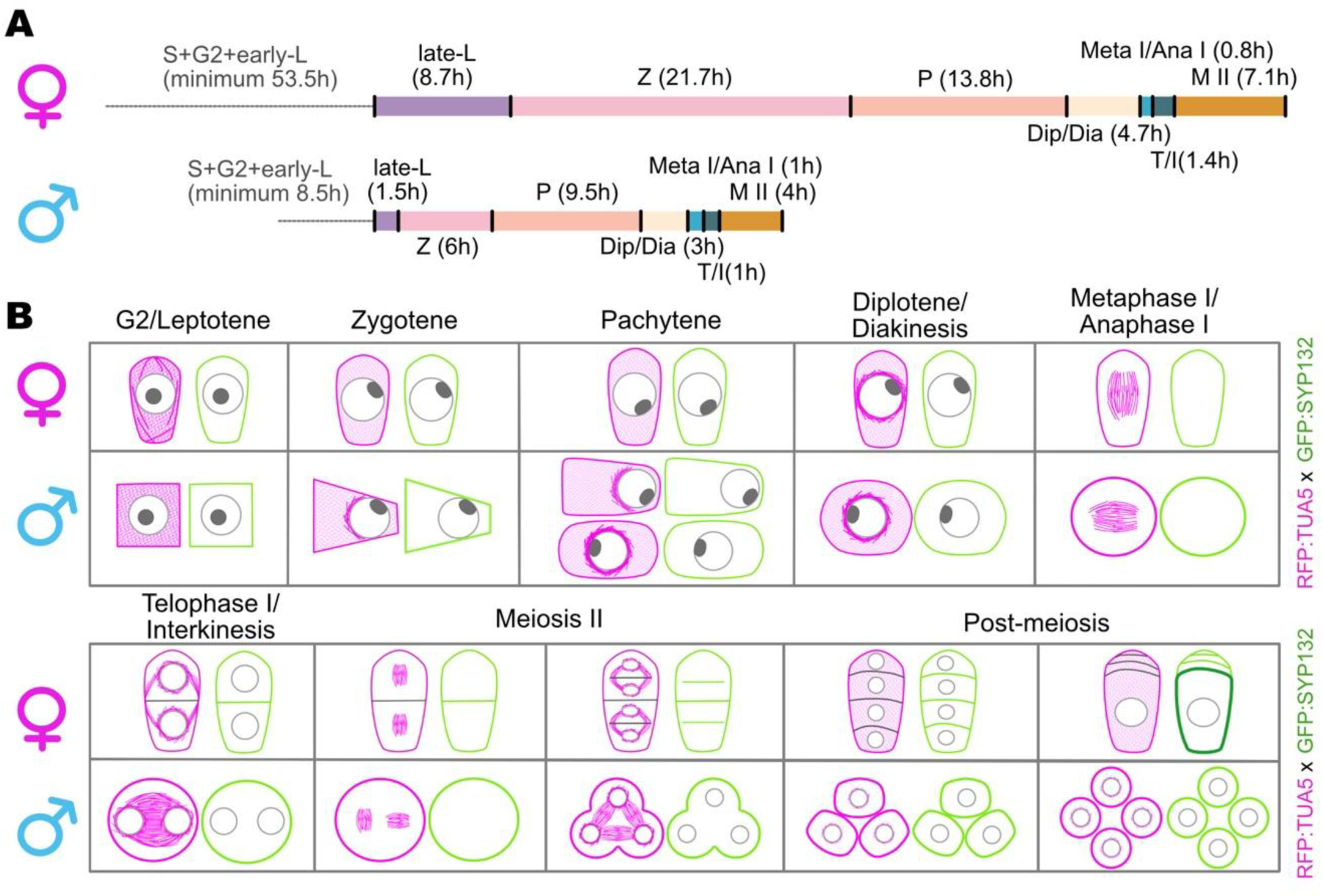
Comparison of female and male meiosis. (A) Comparison of meiotic timelines for female and male meiosis. The female meiotic timeline is based on average durations between defined landmarks as determined in Figure 4C, while durations for male meiosis are taken from Prusicki et al. (2019). S = S-phase; L = Leptotene; Z = Zygotene; P = Pachytene; Dip/Dia = Diplotene and Diakinesis; Meta I/Ana I = Metaphase I and Anaphase I; T/I = Telophase and Interkinesis; M II = Meiosis II. Durations are given in hours. **(B)** Comparison of key meiotic features between female and male meiosis.

Furthermore, late leptotene to diakinesis is also substantially longer in female versus male meiosis (approximately 45h versus 20h). In detail, leptotene in female meiosis is five times longer than in male meiosis, zygotene three times longer, and pachytene, diplotene and diakinesis are each one and a half times longer (Figure 7A). Interestingly, the relative timings are also different and zygotene is much longer than pachytene in female meiosis while in male meiosis, pachytene is longer than zygotene. This suggests that chromosome pairing and synapsis progress much more slowly in female meiosis than in male meiosis. Notably, the chromosome axis is shorter in female meiosis than in male.^6^ This difference has also been found to be associated with the observed heterochiasmy in Arabidopsis.^45^ However, it is currently difficult to determine whether a short chromosome axis is due to the additional time spent in early prophase or whether a short axis requires more time to accommodate synapsis, hence leading to a prolonged prophase. However, both processes could also be controlled independently and/or be controlled by the same upstream regulator(s).

Notably, female meiosis appears to have lower levels of CYCLIN-DEPENDENT KINASE A;1 (CDKA;1) activity, one of the major drivers of meiotic progression.^20,24,46,47^ Furthermore, a modest reduction of CDKA;1 activity on the male side slows down meiosis and triggers cytokinesis after meiosis I. Notably, both cells then enter a second meiotic division completed by a second cytokinesis.^20^ Cytokinesis after the first meiotic division represents the WT situation on the female side, as also visualised here by the SYP132 reporter, further arguing for overall lower levels of CDKA;1 activity during female meiosis. Given that many meiotic regulators contain CDKA;1 consensus phosphorylation sites and might be targets of CDK regulation,^47^ the slower progression on the female side may be the outcome of lower levels of CDKA;1 activity. This would match the observation in mitotically dividing cells where a reduction of CDK activity results in a much delayed progression through the cell cycle.^48,49^ Another difference between female and male meiosis is the shape of the meiocyte. While on the male side, the form of the meiocyte changes during progression through meiosis in the order of rectangular, trapezoidal, oval and circular ^13^, the MMC keeps the same club-shape appearance from the beginning to the end of meiosis.

Whereas the size of the nucleus in both female and male meiocytes consistently increases to a similar extent, we found that the shape and positioning of the nucleus in the MMC differ from those in the male meiocytes. In male meiosis, the nucleus starts in the middle of the meiocytes, moves to one side of the cell at zygotene, and then moves back to the center of the meiocyte at the late pachytene.^13^ In female meiosis, the nucleus, with the exception of small fluctuations, remains in the middle of the meiocyte during prophase I (Figure 7B). In fact, we found several microtubule cables emanating from the side of the cell that apparently anchored the position of the nucleus in the MMC. This anchoring might be also the reason why the wrinkled appearance of the nucleus changes to a spherical shape.

The reason for the positioning is unclear, but it could possibly be linked to the polarity of the MMC (club-shaped) and a division pattern that results in the specification of the most basal of the four meiotic products to become the functional megaspore. With that respect, it is interesting to note that the selection of the most basal cell of the four meiotic products is most likely controlled by cues within the MMC or between them rather than through an external signal such as a hormonal gradient. This conclusion is derived from the observation that in the case of multiple mutants in *KRPs* or in *rbr1* mutants where multiple MMCs are produced, it is still the most basal cell that is selected in each array of meiotic products, independent of their positioning in the ovule primordium.^25^ In contrast, the male meiocytes appear to divide with random division plane orientation and all four meiotic products survive.

However, the nucleolus within the nucleus behaves the same way during both female and male meiotic progression. It only moves from the center to the side during the leptotene stage where it stays until it dissolves in diakinesis. The nucleolus movement was recently found to be linked to the release of the centromeres from the nuclear envelope through the degradation of the nuclear lamina-like CROWN proteins.^41^ This release is crucial for the rapid chromosome movements seen in early prophase of Arabidopsis and other plant species, such as maize.^16,50^ Along with the release of the centromeres from the nuclear envelope, telomeres become detached from the nucleolus and cluster in a restricted region of the nuclear envelope, forming the chromosome bouquet. The chromosomal movement is driven by external forces transmitted by the LINC complex, with SUN1 being one of its components.^17,31,38^ In zygotene of male meiosis, SUN1 is assembled at the inner nuclear envelope at the side of the nucleus that is closest to the cell wall and it displays a crescent like localization. In contrast, the SUN1 domain is highly dynamic in the MMC (Figure 3A,B).

Remarkably, a very prominent microtubule structure, phrased the “half moon,” is absent in female meiosis. The half moon assembles in male meiosis at the side of the nucleus, which faces the cytoplasm at the time when the nucleolus is already placed at the nuclear envelope but moves along the nuclear envelope (Figure 7B). Thus, the half-moon structure appears not to be linked to chromosome movement, at least not primarily, but is rather connected to the movement of the nucleus to one side of the cell in male meiocytes. Furthermore, the microtubule cage on that side of the nucleus appears to hinder the LINC complex from entering this domain, restricting SUN1 to the other side of the nucleus.^38^ Whether this positioning and/or restriction of the LINC complex promotes a faster and/or more efficient chromosome pairing process than in a centrally located nucleus needs to be determined. Clearly, these findings present only the beginning of an understanding of the relevance of microtubule dynamics.

The rapid chromosome movement in zygotene and pachytene is very much slowed down, almost to a complete halt in the diplotene of both female and male meiosis (Movie S5). This arrest coincides with the reappearing of CROWN proteins in the male and possibly a re- anchoring of the centromeres to the nuclear envelope.^17,41^ Whether CROWN proteins also reappear in the MMC at that time point needs to be determined in future.

The arrest of chromosome movement in both female and male meiosis is preceded by a full- moon-like arrangement of microtubules around the nucleus, here described as the perinuclear ring. However the duration of the appearance of this perinuclear ring in female meiosis is much shorter than in males. While it spans in males almost half of pachytene and lasts until the nuclear envelope breaks down, it is only visible from diplotene onwards in females (Figure 7B). Thus, this structure appears to be not, at least not fundamentally necessary for pachytene, e.g., the disassembly of the SC. Instead, the full moon, based on its coinciding with the repositioning of the nucleus to the center of male meiocyte, could play a role in nuclear movement, i.e., generating a pushing force on the other side of the already existing half moon and by that producing a full moon structure.

Taken together, our imaging system revealed many and unexpected cytological differences between female and male meiosis. This comparison helps to refine the possible role of the observed structures, e.g., the half-moon arrangement in the male is likely associated with nuclear positioning rather than with a direct role in chromosome movement. At the same time, this analysis provides criteria to analyze and understand the molecular underpinnings of meiosis in both sexes.

### Commitment to meiosis

Here, we applied our imaging system to analyze meiosis in the quadruple mutant *krp3 krp4 krp6 krp7*, in which a designated female meiocyte undergoes several mitotic divisions before entering meiosis.^25^ This allowed us to approach the questions when a meiocyte is committed to meiosis. On the one hand, this is a fundamental question during germline development. On the other hand, addressing this question has the potential to shed light on diplosporous plants. Diplospory is a form of apomixis in which the embryo sac originates from an unreduced MMC that aborts the normal meiotic divisions.^51^ As a result, the resulting embryo sac retains the somatic chromosome number of the parental plant. This process can lead to the formation of clonal offspring, as the genetic material is passed on without meiotic recombination or reduction.

Our analysis revealed that a meiotic division can be converted to a mitotic division until late into prophase, i.e., at least until the chromosome axis is formed and ASY1 becomes associated with the axis. Notably, we did not see mitotic division after an SC was formed. However, we can currently not say whether the mitosis promoting activity was just diminishing at that moment and hence not sufficient any more to trigger mitosis or whether the formation of the SC presents the point-of-no-return for a meiotic division. Even if the appearance of the SC marks a fixed meiotic division, it remains unclear, whether this is directly dependent on the SC or whether the SC coincides with a factor that cements meiosis.

A conversion of a meiotic division to a mitotic division has also been observed in yeast and is named return-to-growth.^52,53^ This conversion is possible till prometaphase, which has been termed the meiotic commitment point.^54–56^ With that respect, female meiosis in Arabidopsis shares some aspects with yeast, i.e., that mitosis can be induced till relatively late in a meiotic program. A striking difference, however, is that Arabidopsis meiocytes maintain their developmental fate and one or more mitotic divisions do not overwrite the programming to become a meiocyte. Conversely, the repression of mitotic genes is key to meiotic entry in yeast. However, as we and others have shown here and earlier, the situation in plants is different and additional factors are needed to promote meiosis in plants ^45^. Thus, it seems likely that apomictic plants not simply express a mitotic regulator but that other and early acting genes have to be adjusted to an asexual reproduction modus, likely involving an epigenetic reprogramming of developmental fate.^57^ The here-established live-cell imaging system for female meiosis can now serve as a guiding system for future studies that are needed to understand the switch between sexual and asexual reproduction.

## Methods

### Plant material and growth conditions

The Arabidopsis Thaliana plants used in this study were all Columbia (Col-0) background. All the seeds were first surface-sterilized with chlorine gas, then stored in a 4°C fridge overnight. Afterwards, seeds were sown on the 0.7% plant agar plate with half-strength Murashige and Skoog (MS) and 1% sucrose. Antibiotics were added to the plate for selection when required. Then leave the plates with seeds in growth chambers at 21°C for 16h of light and 18°C for 8h of dark cycle for germination. Ten days later, seedlings were transferred to soil and grown in the growth chamber under the same long-day conditions as above, with 70% humidity.

KRP T-DNA insertion lines *krp3* (At5g48820, WsDSLox49707H), *krp4* (At2g32710, Sail_248_B06), *krp6* (At3g19150, Sail_548_B03), *krp7* (At1g49620, GK_841D12), and the triple mutant *krp4krp6krp7* have been described previously.^25,58^ The reporter lines KINGBIRD2 (*PROREC8:REC8:GFPxPRORPS5ATagRFP:TUA5*), *PRORPS5A:TagRFP:TUA5*, *PROASY1:ASY1:TagRFP*, *PROZYP1b:ZYP1b:GFP*, *PROASY3:ASY3:GFP*, *PROSYP132GFP:SYP132*, *PRORPS5A:TagRFP:TUB4*, and *PROSUN1:SUN1:GFP* were described previously.^13,20,23,24,30^ The classical floral dip method was employed for *Arabidopsis thaliana* transformation. All genotypes were confirmed by polymerase chain reaction (PCR) and/or antibiotic selection. All crosses were generated by emasculating the female parent 1 day before anthesis and hand-pollinating 1 to 2 days later.

### Statistical methods

Meiotic phase-specific parametric models for censored time-to-event data were used to estimate median duration. If the exact start and end of a phase are observed (due to the capturing of the flanking landmarks), the respective meiocyte is assumed to have an event, i.e., transition from one phase to the next, and the event time is estimated by the difference between the start and end of this phase. If neither the start nor the end of the phase or only one of both landmarks is observed, the meiocyte is modeled as right-censored. In this case, the censoring time is the observed time. In contrast to De Jaeger-Braet et al. (2021), this sample does not include interval-censored data, as no meiocytes move out of the focal plane and in again. All models include only the intercept and a clustered sandwich estimator of variance to account for the clustered structure of meiocytes within different plants. They differ regarding their underlying distribution, which is chosen based on the Akaike Information Criterion (AIC). Out of the possible distributions (exponential, Gompertz, log-logistic, Weibull, and log-normal), Gompertz was chosen as underlying distribution for the parametric model for stages early leptotene, zygotene, anaphase I and metaphase II, Weibull for stages late leptoene, telophase I / cytokinesis I and telophase II / cytokinesis II, log-normal for stages pachytene, diplotene, interkinesis and tetrads, and loglogistic for stages diakinesis, metaphase I and anaphase II.

Predicted marginal median times and their 95% confidence interval are presented. As this is an exploratory study, the analyses are not adjusted for multiple testing. STATA SE version 17.0 was used for the statistical analysis.

### Live imaging of meiotic division

Flowers of 0.7-1.7 mm were isolated and prepared as shown in the results section. Live-cell imaging setup of female meiosis in Arabidopsis. Within the same ovary, more than one ovule can be imaged. Up to 8 opened ovaries were positioned in the same petri dish and cultured in an agar plate with half-strength Murashige and Skoog (MS) and 1% sucrose. Time-lapse image acquisition was performed using a Zeiss LSM 880 confocal laser scanning microscope, with each time point comprising a Z-stack of 5 or 7 optical sections, spaced 2 to 3 μm apart. The interval time was adjusted from 3 minutes to 10 minutes, depending on the developmental stage and physiological condition of the specimen, in order to balance temporal resolution and phototoxicity. The W-plan Apochromat 40X/1.0 DIC objective was used for image acquisition, which enabled the acquisition of clear, brightfield images for cell shape determination and nucleolus position. Fluorescent signals were captured from three distinct channels. GFP was excited using a 488 nm laser, and emission was collected in the 498-550 nm range. TagRFP was excited at 561 nm, with emission detected between 578-650 nm. Additionally, autofluorescence from chloroplasts was visualized using the same 488 nm excitation wavelength, but with signal collection shifted to the 680-750 nm range to isolate far-red emission. The sample chamber during imaging was stabilized at 21°C to support optimal conditions during long-term observation.

### Image processing

The time-lapse datasets were initially transformed into sequential image stacks representing temporal progression. For each time point, the optimal focal plane was selected manually using aMPkit (unpublished), a software that can review the multi-dimensional data and then export different combinations of xyzt dimension image sequences and save as Tagged Image File Format (TIFF) to preserve uncompressed pixel data. Image drift was corrected by the Stack Reg plugin from Fiji (ImageJ, version 2.16, https://imagej.net/software/fiji).

### Nucleus size measurement

Nuclear diameter was determined by measuring the distance between two parallel lines positioned tangentially to the nucleus using the distance measurement module in NIS-Elements software (version 5.42.02). To account for potential underestimation due to spherical shape and variations in focal plane positioning, three independent measurements were performed per nucleus. The maximum recorded value from these three measurements was selected as the representative diameter to determine the average diameter of leptotene, zygotene, pachytene, and diplotene, ensuring capture of the real equatorial dimension. Statistical comparisons of nucleus diameters between female and male meiocytes at different stages were performed using an independent samples t-test.

### Fluorescence intensity measurement

Fluorescence intensity was measured using Fiji. For each time point and Z-stack, regions of interest (ROIs) corresponding to the nucleus and cytoplasm were manually defined based on the brightfield channel. The cytoplasmic ROI was generated by applying the XOR function to subtract the nuclear region from the boundary. Similarly, the nucleolus was excluded from nuclear measurements by defining a smaller inner ROI representing the nucleolus and using the XOR function to isolate the nucleoplasm. Fluorescence intensities were measured across all time points in the relevant fluorescence channel, and the mean grey values were used for quantification.

### Declaration of generative AI and AI-assisted technologies in the writing process

During the preparation of this work, the authors used ChatGPT in order to correct the language use. After using this tool/service, the authors reviewed and edited the content as needed and take full responsibility for the content of the publication.

## Acknowlegments

We would like to thank Dr. Anika Bucholz (UKE, Hamburg) for her support in the data analysis. We are further grateful to Dr. Maren Heese (University of Hamburg) for critical reading and constructive comments on the manuscript. BH was supported by a fellowship of the Elisabeth Appuhn-Stiftung to complete her doctoral training. This work was supported by DFG grants SCHN 736/21-1 and SCHN 736/15-1 to AS.

## Author contributions

B.H., Conceptualization, Investigation, Visualization, Writing - original draft, Writing - review and editing; M.P., Conceptualization, Investigation, Visualization; K.S., Formal analysis, Software; Y.W., Investigation; A.S., Conceptualization, Funding acquisition, Project administration, Resources, Supervision, Writing - original draft, Writing - review and editing.

## Supplemental information

Document S1. Figure S1 and Table S1

Movie S1. Microtubule behavior and REC8 loading. Live cell imaging of REC8:GFP x RFP:TUA5 was performed in a female meiocyte. Microtubule (magenta) bundles extend from the spindle pole during REC8 (green) loading, and REC8 signals become visible at 30 minutes. The movie starts before REC8 loading and runs for 1060 minutes with scanning intervals of 10 minutes. Scale bar: 10 μm.

Movie S2. Chromosome and microtubule behavior from late pachytene to telophase II. Live cell imaging of REC8:GFP x RFP:TUA5 was performed in a female meiocyte. REC8 signals become dimmer over time. The first NEB at 600 min and the second at 720 min. The movie starts at late pachytene and runs for 820 minutes with scanning intervals of 10 minutes. Scale bar: 10 μm.

Movie S3. Chromosome dynamics and nucleolus movement in early prophase I. Live cell imaging of REC8:GFP was performed in a female meiocyte. REC8 signal intensifies and gradually forms thin threads over time, while the nucleolus (absent of REC8 signal) migrates from the center to the nuclear envelope. Thin threads further synapse into thick threads. The movie runs for 1160 minutes with scanning intervals of 10 minutes. Scale bar: 10 μm.

Movie S4. Visualization of chromosome synapsis by the loading of ZYP1b and the removal of ASY1. Live cell imaging of ZYP1b:GFP x ASY1:RFP was performed in a female meiocyte. The movie starts at zygotene and runs for 1160 minutes with scanning intervals of 10 minutes. Scale bar: 10 μm.

Movie S5. Visualization of chromosome rapid movement during prophase I. Live cell imaging of ASY3:GFP x RFP:TUA5 was performed in a female meiocyte. The movie starts at late leptotene and runs for 950 minutes with scanning intervals of 10 minutes. Scale bar: 10 μm.

Movie S6. Rapid prophase chromosome movements slow down while the microtubules form a perinuclear ring. Live cell imaging of ASY3:GFP x RFP:TUA5 was performed in a female meiocyte. The chromosome movements slow down at 190 min. The movie starts at late pachytene and runs for 460 minutes with scanning intervals of 10 minutes. Scale bar: 10 μm.

Movie S7. Phragmoplast and cell wall formation. Live cell imaging of GFP:SYP132 x RFP:TUA5 was performed in a female meiocyte. The first NEB at 160 min and the second at 290 min. The movie starts at late prophase I and runs for 400 minutes with scanning intervals of 10 minutes. Scale bar: 10 μm.

Movie S8. A presumptive MMC divides mitotically before REC8 loading in *krp3 krp4 krp6 krp7*. Live cell imaging of REC8:GFP x RFP:TUA5 was performed in a female meiocyte of *krp3 krp4 krp6 krp7.* The NEB is at 180 min prior to REC8 loading, resulting in two products without any detectable REC8 signals. The movie runs for 1090 minutes with scanning intervals of 10 minutes. Scale bar: 10 μm.

Movie S9. A REC8 expressing MMC is divided mitotically in *krp3 krp4 krp6 krp7*. Live cell imaging of REC8:GFP x RFP:TUA5 was performed in a female meiocyte of *krp3 krp4 krp6 krp7.* The upper MMC goes through the mitotic division at 320 min, resulting in two products with intensified REC8 signals (green) over time. The movie runs for 780 minutes with scanning intervals of 10 minutes. Scale bar: 10 μm.

Movie S10. Mitotic division happens both before and after ASY1 loading in *krp3 krp4 krp6 krp7.* Live cell imaging of ZYP1b:GFP x ASY1:RFP was performed in a female meiocyte of *krp3 krp4 krp6 krp7.* The movie starts with two MMCs, the upper one without ASY1, while the lower one with ASY1 (magenta) loading over time. The lower MMC divides at 360 min, resulting in two products with stronger ASY1 expression, but one daughter cell lost the ASY1 expression at 2050 min. The upper MMC follows the division at 460 min, with no ASY1 signals in the daughter cells. The movie runs for 2190 minutes with scanning intervals of 10 minutes. Scale bar: 10 μm.

Movie S11. Meiosis division happens as long as ZYP1 is loaded in *krp3 krp4 krp6 krp7.* Live cell imaging of ZYP1b:GFP x ASY1:RFP was performed in a female meiocyte of *krp3 krp4 krp6 krp7.* The movie starts with two MMCs with ASY1 expression (magenta). The upper one gets ZYP1b loading at 430 min, the first NEB at 2070 min and the second at 2190 min. The lower MMC gets ZYP1b loading at 530 min, the first NEB at 2430 min and the second at 2560 min. The movie runs for 2670 minutes with scanning intervals of 10 minutes. Scale bar: 10 μm.

Excel sheet 1. Contiguous representation of meiotic stages from 105 movies. Script 1. Code for time course analysis.

